# Pharmacodynamic modeling of colistin and imipenem against *in vitro Pseudomonas aeruginosa* biofilms

**DOI:** 10.1101/2024.05.28.596312

**Authors:** Yuchen Guo, Jinqiu Yin, Linda B.S. Aulin, Oana Ciofu, Claus Moser, Parth J. Upadhyay, Niels Høiby, Hengzhuang Wang, Tingjie Guo, J. G. Coen van Hasselt

## Abstract

**Introduction:** Antibiotic treatment of chronic biofilm-associated infections can be challenging. Characterization of pharmacokinetic-pharmacodynamic (PK-PD) relationships for biofilm-associated infections may be relevant to inform the design of antibiotic treatment regimens for biofilm-associated infections. To this end, we aim to develop a mathematical PK-PD model for planktonic and biofilm bacterial infections and demonstrate how PK-PD simulations can be used to design optimized dosing schedules, using imipenem and colistin as proof-of-concept examples.

**Methods:** Pharmacodynamic models were developed using time-kill assay data from planktonic and alginate-bead biofilm cultures of *Pseudomonas aeruginosa* exposed to imipenem or colistin. The PD models were coupled to population PK models for plasma and epithelial lining fluid (ELF) to translate PD relationships for clinical dosing schedules and PK-PD indices.

**Results:** The developed models incorporated sensitive and resistant bacterial subpopulations and were able to adequately capture the observed time-kill data. Simulation studies identified differences in suppression of bacterial growth dynamics for multiple clinical intravenous and inhalation-based treatment regimens and were used to infer biofilm-specific PK-PD indices associated with ELF target site concentrations.

**Conclusion:** In conclusion, we demonstrate the utility of mathematical modeling for the characterization of PK-PD relationships underlying time-kill kinetic profiles in biofilm-associated infections and their utility in translating experimental findings to inform the optimization of clinical dosing schedules.

## INTRODUCTION

Chronic lung infections associated with cystic fibrosis (CF) are typically associated with bacterial biofilms and respond poorly to antibiotic therapy [1–3]. Patients with chronic CF lung infections may receive long-term antibiotic therapy including daily nebulized antibiotic treatment and systemic antibiotic treatment during acute exacerbations [4, 5]. Biofilm-associated pathogens often show reduced antibiotic sensitivity compared to their planktonic form, mediated by several mechanisms [6]. In addition, the antimicrobial target-site concentrations may differ significantly from plasma concentrations, i.e., the lungs in case of chronic CF lung infections [7, 8]. There is a need to further optimize antibiotic dosing schedules for the treatment of biofilm-associated chronic lung infections in CF patients.

A rational treatment design for biofilm-associated bacterial infections requires information on the antibiotic concentration-time profile at the site of infection (pharmacokinetics, PK) and the observed relationship between drug exposure and response of bacterial pathogens (pharmacodynamics, PD). In terms of PD, different mechanisms contribute to the decreased susceptibility of biofilm bacteria [9]. For example, the formation of extracellular matrix protects the inside bacteria from the attack of immune system and poses a diffusion barrier against antibiotics. In addition, bacterial pathogens may develop resistance, i.e., resilience against antibiotic treatment mediated through transient adaptation or non-transient genetic mutations.

Antimicrobial PK-PD relationships can be characterized using experimental *in vitro* and *in vivo* models. Although static *in vitro* assays such as MIC or MBIC are useful to obtain quick insight into antimicrobial sensitivity, they are evaluated at a single time point, for example, 24 h, and do not provide information on dynamic responses such as the emergence of transient or non-transient antimicrobial resistance [10]. In contrast, *In vitro* and *in vivo* time-kill assays enable characterization of the time course of bacterial response to antimicrobial agents [11–13], providing essential information about pathogen-associated PD relationships.

Mathematical mechanism-based PD models are useful tools in quantitatively characterizing the bacterial growth and kill dynamics determined by time-kill assays. Such mechanism-based PD models support systematic testing of hypotheses that may explain observed pharmacodynamic responses with respect to delays (*e*.*g*., due to drug diffusion), differences in growth rates of bacterial subpopulations, and the shape of concentration-effect relationships [14]. More importantly, when PD models are coupled to population PK models that predict antimicrobial concentration-time profiles in patients [13, 15, 16], the efficacy of clinical dosing schedules can be evaluated to assess alternative optimal dosing regimens. Most antimicrobial PK-PD models have focused on planktonic bacterial pathogens, lacking attention to biofilm-associated pathogens [17]. To optimize the dosing schedules for biofilm-associated infections, the characterization of PK-PD relationships for biofilm-associated infections is necessary.

Here, we aim to demonstrate the utility of mathematical PK-PD modeling for the analysis of experimental biofilm time-kill studies to ultimately guide the optimization of dosing schedules for biofilm-associated infections. We focus on imipenem and colistin for the treatment of the CF-associated pathogen *P. aeruginosa*, as proof of concept. We specifically aim to (1) develop PD models for imipenem and colistin using data generated from *in vitro* time-kill studies in planktonic and alginate-bead biofilm experiments [18, 19] and (2) pair the developed models to population PK models for plasma and lung concentrations to explore and evaluate dosing schedules and PK-PD targets for biofilm-associated infections, compared to planktonic infections.

## METHODS

### Time-kill studies

Previously published time-kill studies of *P. aeruginosa* PAO1 planktonic and alginate bead biofilm cultures exposed to imipenem or colistin were used for mathematical model development [18, 19]. Briefly, the inoculum for both planktonic and alginate bead experiments was 10^6^ CFU/mL in lysogeny broth medium. Beads (50-100 µm) were produced by embedding *P. aeruginosa* in seaweed alginate [19]. For colistin, both planktonic and alginate bead biofilm cultures were exposed to colistin at concentrations of 0-256 mg/L for 24 hours. For imipenem, planktonic cultures were tested against imipenem at concentrations of 0-32 mg/L for 24 hours, while for the alginate bead biofilm experiments, additional concentrations up to 2048 mg/L were included (**Table S1**). The studied concentrations covered a relatively wide efficacy range. Samples were taken for CFU quantification at 0, 1, 2, 4, 8, 12 and 24 hours post antibiotic exposure.

### Mathematical model development

Ordinary differential equation (ODE)-based compartmental models were developed to describe the bacterial growth and kill dynamics in planktonic and biofilm cultures. The models included subpopulations of sensitive (S) and resistant (R) bacteria, where resistant bacteria were assumed to reflect a bacterial subpopulation with reduced sensitivity. Colony forming units (CFU) data were log_10_-transformed prior to the analysis. Models for planktonic and biofilm bacteria were developed separately. Log-transformed predictions were used to estimate the parameters that maximized the log-likelihood.

We incorporated natural growth kinetics, the net growth of bacteria in absence of antibiotic, for planktonic and biofilm cultures. A capacity-limited growth model was used (Eq. 1), including parameters for the maximum bacterial density (*B*_*max*_), and a first-order net growth rate (*k*_*gs*_), with a starting bacterial density (CFU/mL) of *B*_*0*_.

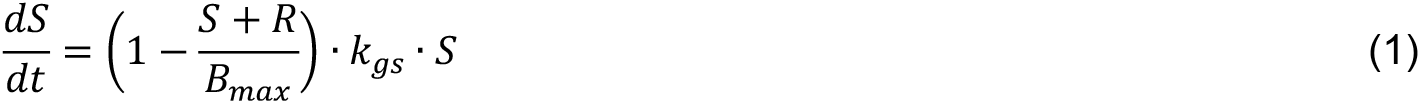

Drug concentration-effect functions evaluated included linear (Eq. 2) and (sigmoid) E_max_ functions (Eq. 3), separately, for each individual subpopulation (i.e., S, R). Antibiotic concentration-effect models (i.e., linear or Emax) were defined as follows:

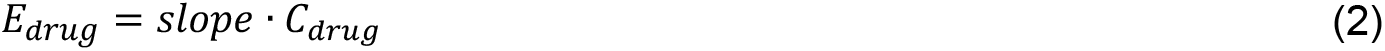

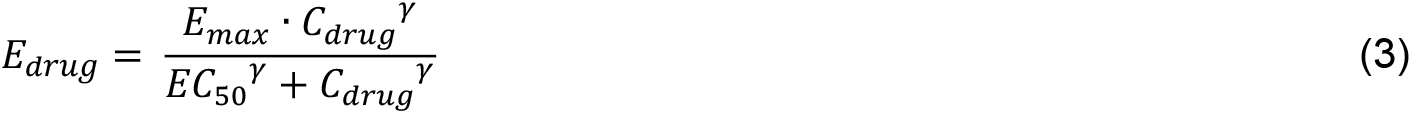

where *slope* is the linear kill rate constant, *E*_*max*_ represents the maximum kill effect, *EC*_50_ indicates the drug concentration at which 50% of the maximum effect is obtained, and γ is the steepness of the concentration-effect relationship factor. Since drug concentration-effect models may vary across drugs, bacterial subpopulations and lifestyles (i.e., planktonic or biofilm), separate drug effect models were considered for each of these conditions.

We investigated the occurrence of an effect delay in biofilm cultures, e.g., which could for example be explained by retarded drug diffusion into the biofilm. Such a delay would account for a possible discrepancy between the experimentally used drug concentration 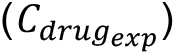 and the effective drug concentration that exerted pharmacodynamic effect on bacteria. The delay was described using a transit model (Eq. 4), with a first-order transit rate constant *k*_*tr*_, including *n* transit compartments. The concentration in the last transit compartment 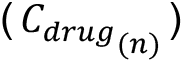 represented the concentration driving the effect. Mean transit time (*MTT*), the average time spent by drugs traveling from the first transit compartment to the last compartment, was calculated with *k*_*tr*_ and *n* (Eq. 5).

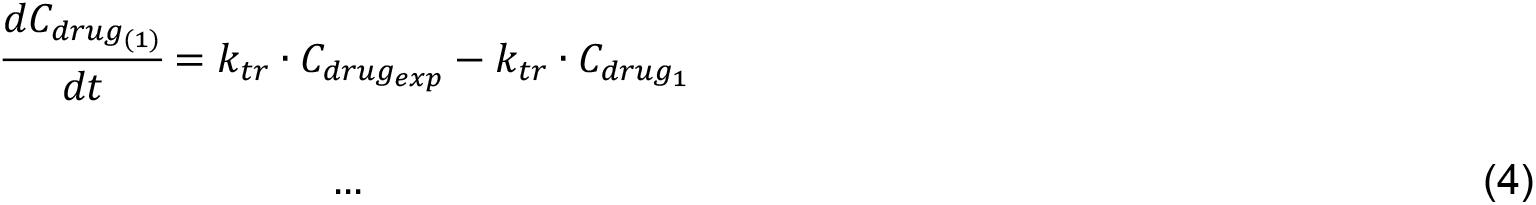

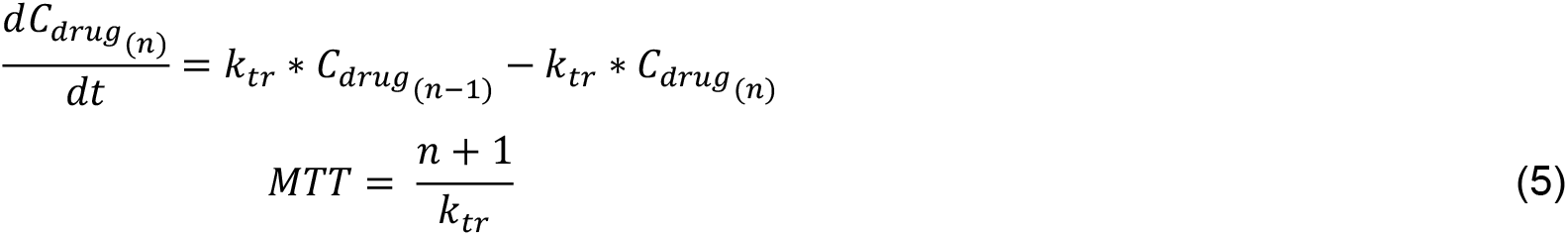

We incorporated a drug-induced transition rate *k*_*sr*_ to describe the transfer of bacteria from S to R state, which only occurred if an antibiotic is present (Eq. 6-7), for both planktonic and biofilm bacteria. Initially, all bacteria were assumed to be in the sensitive (S) state. First-order growth rates for S and R populations were estimated separately. Drug-induced killing effect was described using a first-order rate process for each subpopulation separately.

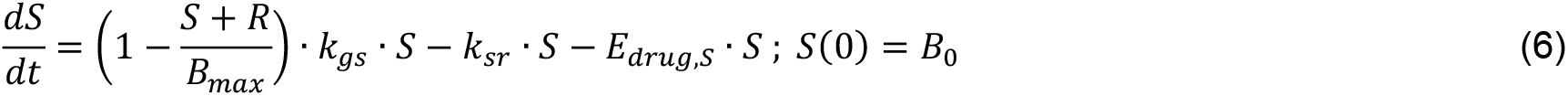

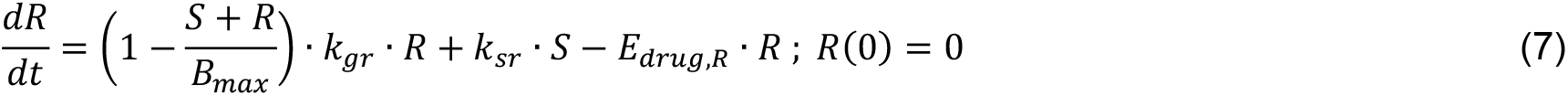

An additive error model for log-transformed data was used to estimate residual unexplained variability. Bacterial counts below the lower limit of quantitation (LLOQ, defined as 10 CFU/mL) were handled using the M3 method [20].

### Sensitivity analysis

To determine the relative importance of model parameters estimated, a sensitivity analysis was performed for each parameter (*p*) in the final model. The local sensitivity *Sens* was evaluated using the relative change in the area under time-CFU curve (AUC) between 0 and 24 hours, in relation to the relative change of parameters (Eq. 8) [21].

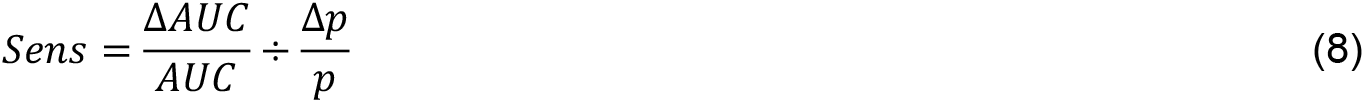

### Dosing regimen simulations

We implemented published population PK models for intravenous and inhaled colistin and intravenous imipenem [22, 23] to predict the antibiotic concentration-time profiles in plasma and epithelial lining fluid (ELF) in the lung. 1000 individuals were simulated to take the inter-individual variability into account. As patient covariates (i.e. body weight and creatinine clearance) acted as significant factors in the prediction of the concentration profiles of imipenem, a virtual population generated from the NHANES copula (https://cocosim.lacdr.leidenuniv.nl/) was used as the input to obtain realistic covariates combinations and simulation results [24, 25]. For colistin, we studied intravenous administration of 160 mg (2 MIU) every 8 hours of colistimethate sodium (CMS), the inactive prodrug of colistin, 720 mg (9 MIU) CMS every 24 h, and inhalation of 160 mg (2 MIU) CMS every 8 hours, consistent with recommended clinical dosing regimens [26]. For imipenem, we simulated clinical tolerable doses [4]: 250, 500 and 1000 mg every 6 hours intravenously. The PK simulations for ELF antibiotic concentrations were linked to our PD models for planktonic and biofilm bacteria, to study the relative difference in bacterial dynamics under different dosing schedules for planktonic and biofilm-associated infections. Protein concentrations in ELF were considered negligible.

### PK-PD target analyses

To identify PK-PD indices for colistin and imipenem relating to planktonic and biofilm bacteria, we simulated extensive dose fractionation studies, using a wide dose range for a duration of 24 hours, similar to the method used by a previous study [27]. For each dosing schedule, we computed the PK-PD indices, including the maximum ELF concentration of drug over the minimum inhibitory concentration (C_max_/MIC) and over the minimum biofilm inhibitory concentration (C_max_/MBIC), area under the concentration-time curve for drug over the MIC (AUC/MIC) and over the MBIC (AUC/MBIC), and the fraction of time when the concentration was above the MIC (f_T>MIC_) and above the MBIC (f_T>MBIC_). The PK profiles were used to predict the treatment response in planktonic and biofilm bacterial infections using the established PD models. For each bacterial lifestyle against each drug, we regressed PK-PD indices against the change of bacterial density (log10 CFU/mL) after 24 hours of treatment using a sigmoidal E_max_ equation, and the fit was evaluated by calculating the R^2^ value, to select the PK-PD indices that could best predict the killing effect after 24 hours (e.g.-1 and - 2 log10 kill).

### Software and model selection

Model development was performed using non-linear mixed fixed effects modeling software NONMEM 7.5. Graphical visualizations, simulations and interpretations were performed using R 4.3.2. Model selection was guided by successful minimization, successful covariance step, objective function value (OFV, 5% significance level), Akaike information criterion (AIC), precision of parameters’ estimates (relative standard errors < 30%), goodness-of-fit plots, and visual predictive checks [28].

## RESULTS

### Mathematical model development

Both colistin and imipenem showed a concentration-dependent killing effect against planktonic and biofilm-embedded *P. aeruginosa*. Regrowth was observed after approximately 4 hours in both models when exposed to colistin and 4-8 hours when exposed to imipenem (**Figure S1**). The time-kill kinetics of planktonic and alginate bead biofilm cultures were described separately using pharmacodynamic models for colistin (**Figure 1A**) and imipenem (**Figure 1B**). The models adequately captured the observed bacterial growth- and kill-profiles (**Figure 2**). Model parameters were estimated precisely, with relative standard errors ≤ 20% (**Table 1**).

**Table 1.** Parameter estimates of the pharmacodynamic model for imipenem and colistin for *in vitro* planktonic and biofilm time-kill assays.

| Parameter | Description | Units | Colistin (RSE%) |  | Imipenem (RSE%) |  |
| --- | --- | --- | --- | --- | --- | --- |
|  |  |  | Planktonic | Biofilm | Planktonic | Biofilm |
| $k_{gs}$ | Maximal growth rate of sensitive bacteria | $h^{-1}$ | 0.807 (6) | 0.615 (7) | 0.944 (3) | 0.441 (11) |
| $k_{gr}$ | Maximal growth rate of resistant bacteria | $h^{-1}$ | 3.71 (21) | 3.85 (2) | 1.76 (4) | 0.0614 (7) |
| $B_0$ | Baseline bacterial count | $\log_{10}$ CFU/mL | 5.81 (1) | 5.45 (3) | 5.58 (2) | 6.45 (2) |
| $B_{max}$ | Bacterial count in stationary phase | $\log_{10}$ CFU/mL | 9.35 (1) | 8.85 (2) | 9.42 (1) | 9.38 (4) |
| $k_{sr}$ | Transfer rate from sensitive bacteria to resistant bacteria | $-\log_{10} h^{-1}$ | 3.96 (16) | 4.27 (2) | 3.74 (1) | 3.11 (9) |
| $Slope_s$ | Linear coefficient for drug effect in sensitive bacteria | L/mg*h | 1.29 (3) | 1.21 (7) | - | 0.149 (11) |
| $Slope_r$ | Linear coefficient for drug effect in resistant bacteria | L/mg*h | 0.86 (20) | - | 0.279 (5) | 0.0008 (20) |
| $E_{max_s}$ | The maximal achievable kill rate constant for sensitive bacteria | $h^{-1}$ | - | | 9.28 (1) | - |
| $EC_{50s}$ | Concentration that results in 50% of $E_{max}$ for sensitive bacteria | mg/L | - | | 19.1 (1) | - |
| $Hill_s$ | Hill coefficient for sensitive bacteria | | - | | 0.285 (4) | - |
| $E_{max_r}$ | The maximal achievable kill rate constant for resistant bacteria | $h^{-1}$ | - | 4.06 (2) | - | - |
| $EC_{50r}$ | Concentration that results in 50% of $E_{max}$ for resistant bacteria | mg/L | - | 0.416 (4) | - | - |
| $Hill_r$ | Hill coefficient for resistant bacteria | | - | 0.399 (3) | - | - |
| $k_{tr}$ | Distribution rate constant between transit compartments | $h^{-1}$ | - | 0.66 (3) | - | 1.19 (8) |
| $N$ | Number of transit compartments | - | - | 1 | - | 3 |
| RES | Additive residual error in bacteria experiment ($\log_{10}$ scale) | - | 0.364 (8) | 0.564 (7) | 0.394 (6) | 0.898 (6) |

RSE, relative standard error; RES, residual variance error.

**Figure 1.**
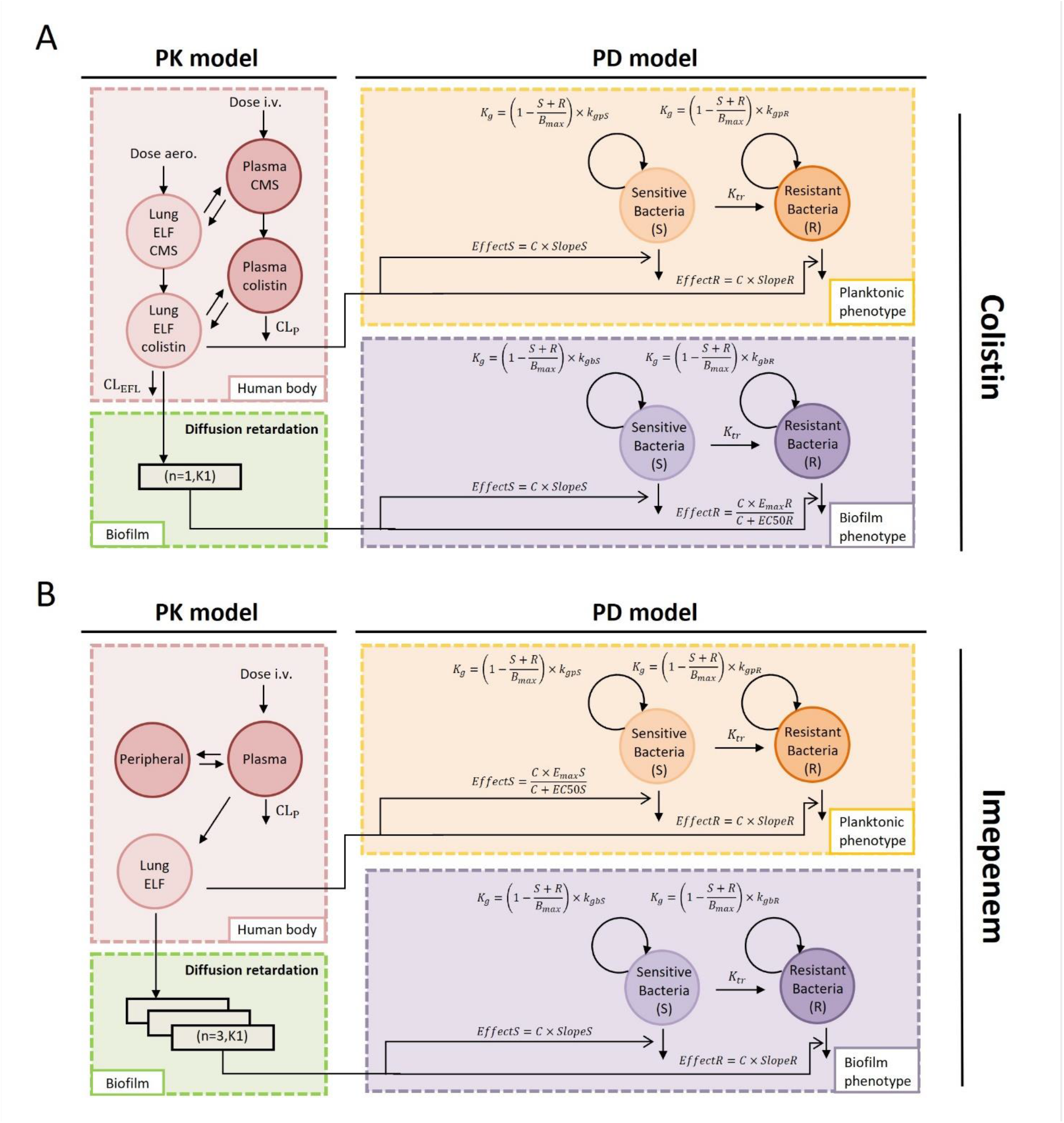
Schematic illustration of the pharmacokinetic-pharmacodynamic model structures. Model structures for colistin and imipenem included the pharmacodynamic models describing the time-kill kinetics in planktonic and alginate bead biofilm models, in combination with clinical pharmacokinetic models for plasma and lung epithelial lining fluid (ELF) compartments. Separate pharmacodynamic models were developed for biofilm and planktonic bacteria cells for colistin and imipenem.

**Figure 2.**
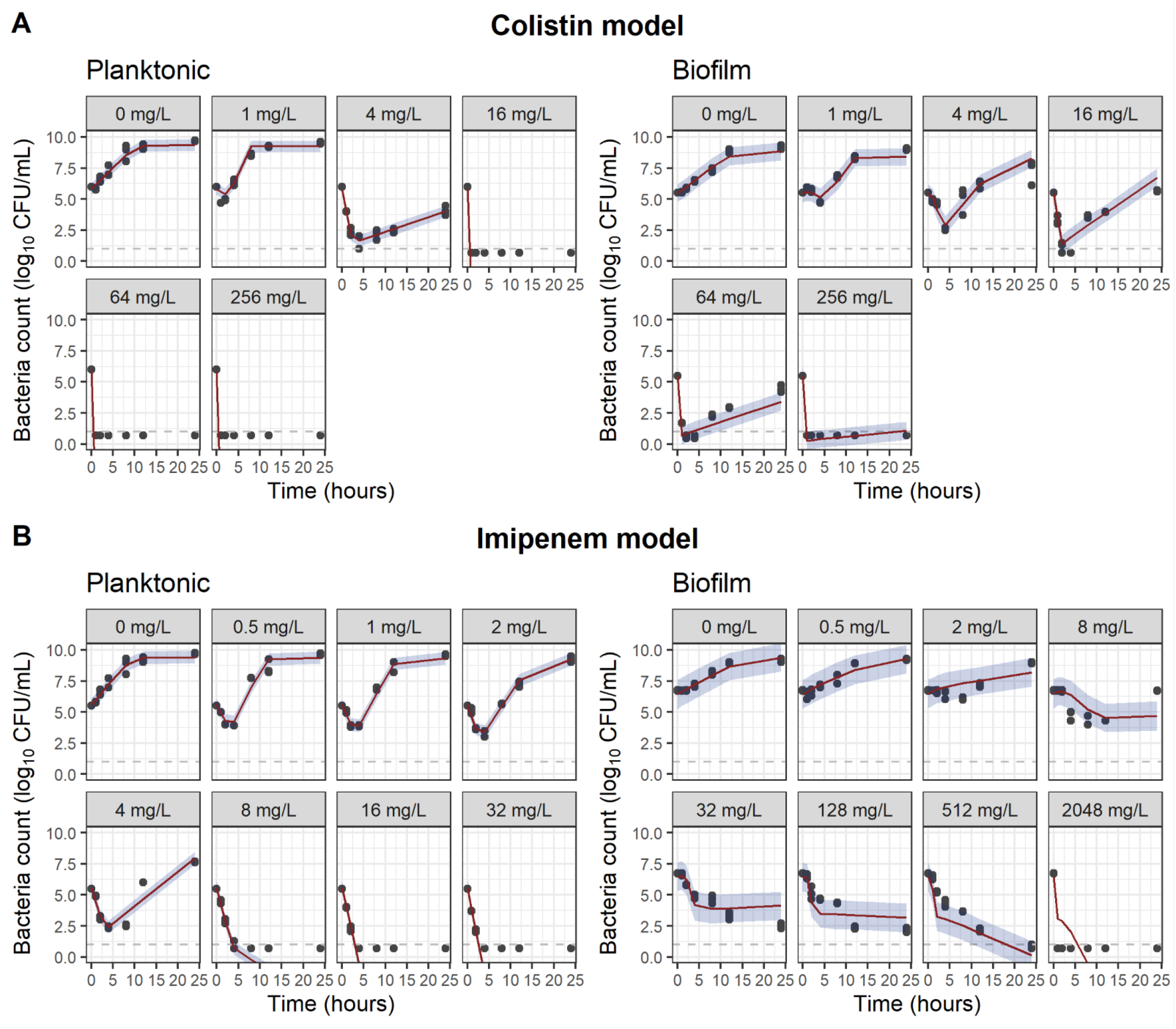
Visual predictive check of the pharmacodynamic models for colistin (A) and imipenem (B). The black points are the observed bacterial count data (Log10 CFU); the red lines represent the median values of model predictions; The blue areas are the 10^th^ to 90^th^ percentile area of the model predictions. Observations below the quantification limit (gray dashed line) were displayed as half of the quantification limit.

Emax models best described the drug concentration-effect relationship for colistin on the biofilm resistant bacteria population and imipenem on the planktonic sensitive bacteria population, while linear models were identified for all other drug effect relationships. *E*_*drug*_ of colistin and imipenem on S and R bacteria populations demonstrated comparable efficacy for planktonic bacteria across drug concentrations and showed less than 0.5 fold differences at the highest tested concentrations (**Figure 3**, left panel). However, significant differences were observed in *E*_*drug*_ between sensitive and resistant biofilm populations, with the difference increasing for higher drug concentrations (**Figure 3**, right panel), suggesting increased resistance in the biofilm population compared to planktonic populations.

**Figure 3.**
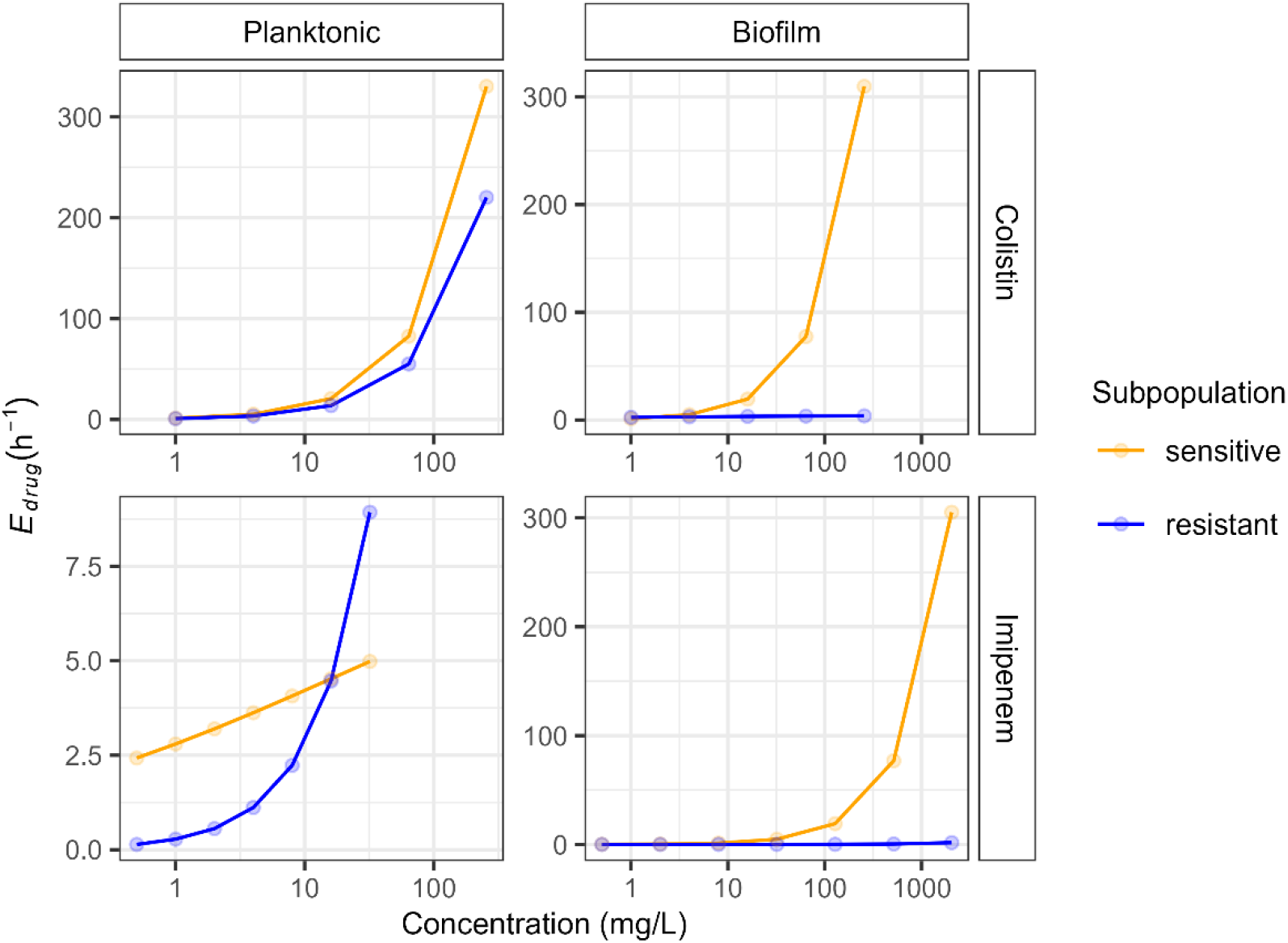
Drug effect comparison. The intermediate variable, *E*_*drug*_, was computed for each drug concentration studied in the time-kill assay using estimated parameters derived from drug concentration-effect models. Orange and blue lines represent exposure-response curves of sensitive and resistant subpopulations, respectively. Points on lines indicate the drug effect at the concentrations studied in time-kill experiments.

For both colistin and imipenem, we estimated the drug-specific rate of diffusion retardation into the biofilm. One transit compartment was implemented for colistin, while three transit compartments were employed for imipenem. We also evaluated models with 0 to 4 transit compartments, and found 1 and 3 transit compartment(s) fitted the colistin and imipenem data best. Parameter estimates (*k*_*tr*_ and *n*) revealed that imipenem exhibited a slightly longer MTT (3.4 hours) compared to colistin (3.0 hours). Local sensitivity analysis identified the most sensitive parameters driving the predicted response for each model, with resistant subpopulation-related parameters (i.e. *slope*_*R*_, *k*_*sr*_, *E*_*max,R*_ and *EC*_50,*R*_) demonstrating wide influence across both studied drugs (**Figure S2**).

### Dosing regimen simulations

We simulated the treatment outcome of standard and adapted dosing schedules of colistin (**Figure 4**) and imipenem (**Figure 5**) against planktonic and biofilm-associated *P. aeruginosa* lung infections, using the PK-PD models (i.e., developed PD models coupled with clinical population PK models).

**Figure 4.**
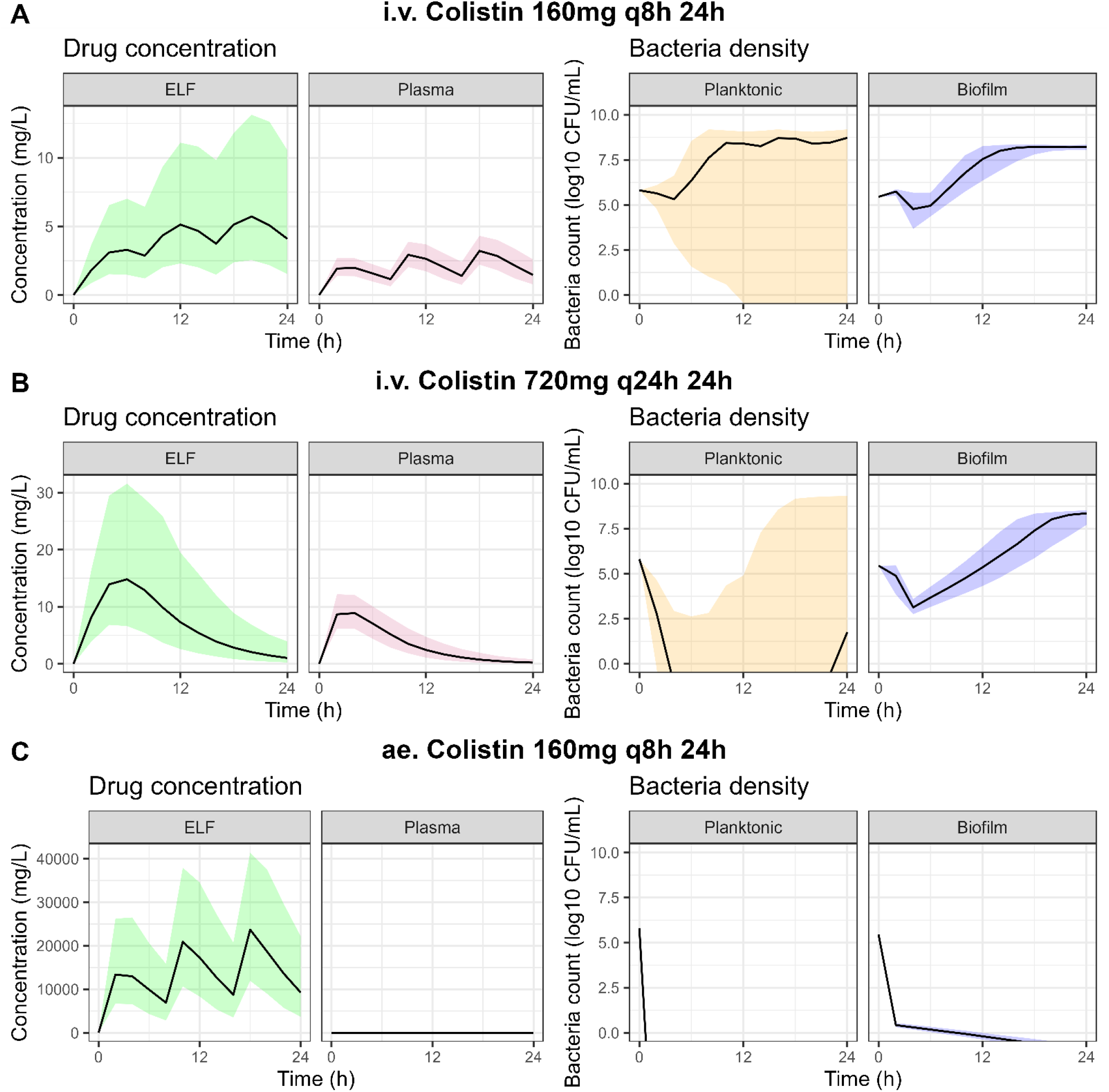
Dosing regimen simulations for colistin. Dosing regimens were simulated using clinical population pharmacokinetic models depicting the median (lines) and 25^th^ and 75^th^ percentiles (shaded areas) of drug concentration and bacteria count versus time. The predicted epithelial lining fluid (ELF) concentration was used as input for the pharmacodynamic models. Dosing regimens simulated included intravenous (i.v.) administration of 160 mg (2 MIU) q 8 h (**A**), 720 mg (9 MIU) q 24 h (**B**), or aerosol (ae.) inhalation of 160 mg (2 MIU) q 8 h (**C**).

**Figure 5.**
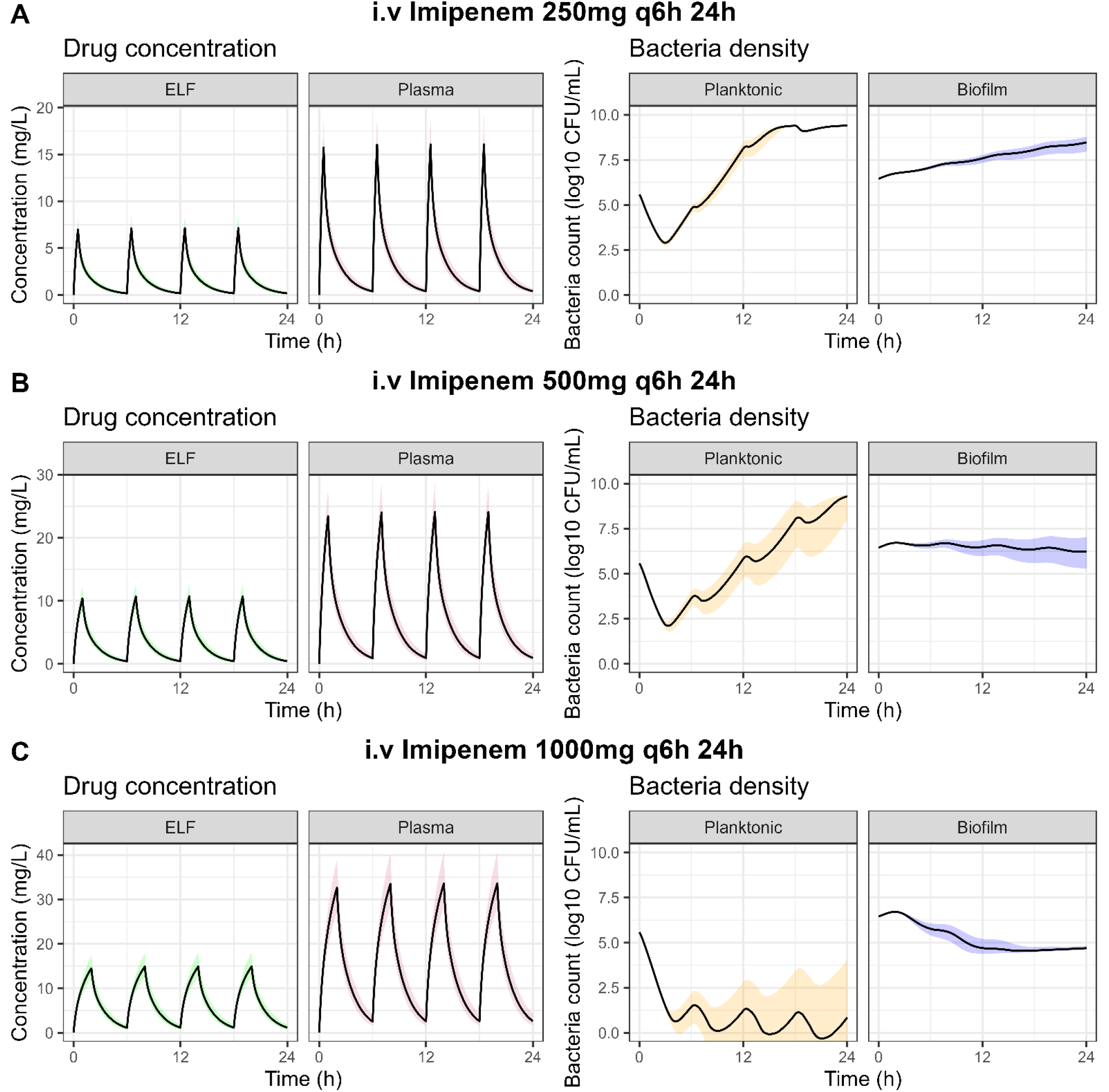
Dosing regimen simulations for imipenem. Dosing regimens were simulated using clinical population pharmacokinetic models depicting the median (lines) and 25 and 75^th^ percentiles (shaded areas) of drug concentration and bacteria count versus time. The predicted epithelial lining fluid (ELF) concentration was used as input for the final pharmacodynamic models. Dosing regimens simulated included intravenous (i.v.) administration of 250 mg (**A**), 500 mg (**B**) or 1000 mg (**C**) every 6 hours.

For colistin, our simulations showed that under the treatment of 160 mg (2 MIU) every 8 hours (**Figure 4A**) and 720 mg (9 MIU) every 24 hours (**Figure 4B**), colistin was insufficient for patients with biofilm infections. An inhalation dose of 160 mg (2 MIU) resulted in a high drug concentration at the ELF, leading to a successful eradication of both planktonic and biofilm bacteria (**Figure 4C**).

For imipenem, a clear dose-dependent killing effect was found. For planktonic bacterial infection, both 250 mg and 500 mg every 6 hours (**Figure 5A, 5B**) could suppress the growth of biofilm cells yet were unable to fully eliminate the infections within 24 hours; 1000 mg every 6 hours (**Figure 5C**) could efficiently kill the bacteria.

### PK-PD indices

We investigated the PK-PD indices for the prediction of treatment response of planktonic and biofilm bacteria infections by performing intravenous dose fractionation studies (**Figure 6**), based on the target site drug concentration in the ELF. We found that AUC/MIC (R^2^=0.995 for colistin; R^2^=0.997 for imipenem) and AUC/MBIC (R^2^=0.956 for colistin; R^2^=0.997 for imipenem) were best correlated with the observed effect for both planktonic and biofilm bacterial infections for both drugs. For planktonic bacteria, we found that for a CFU change from -1 log_10_ to -2 log_10_, the AUC/MIC target for colistin and imipenem increased from 46 to 48, and from 123.7 to 130.6, respectively. For biofilm bacteria, a CFU change of - 1 log_10_ at 24 hours corresponded to 567.1 and 3 for the AUC/MBIC target of colistin and imipenem, respectively. However, no targets of AUC/MBIC were able to be derived for -2 log_10_ CFU change due to insufficient drug effect at tolerable intravenous dosages (**Table 2**). The treatment outcome under simulated dosing regimens aligned with the expected response based on the derived targets. For example, 160 mg (2 MIU) colistin every 8 hours (AUC/MIC = 39) did not achieve the PK-PD targets for planktonic infection and failed to eradicate the bacteria, while 720mg every 24 hours (AUC/MIC = 70) reached the targets and eradicated the bacteria efficiently.

**Table 2.** ELF-based PK-PD target values derived for colistin and imipenem at -1 and -2 log10 unit kill using simulated dose fractionation studies.

|  | Colistin |  | Imipenem |  |
| --- | --- | --- | --- | --- |
|  | Planktonic | Biofilm | Planktonic | Biofilm |
|  | AUC/MIC | AUC/MBIC | AUC/MIC | AUC/MBIC |
|  | (h) | (h) | (h) | (h) |
| **-1 $\log_{10}$** | 46 | 567.1 | 123.7 | 3 |
| **-2 $\log_{10}$** | 48 | - | 130.6 | - |

**Figure 6.**
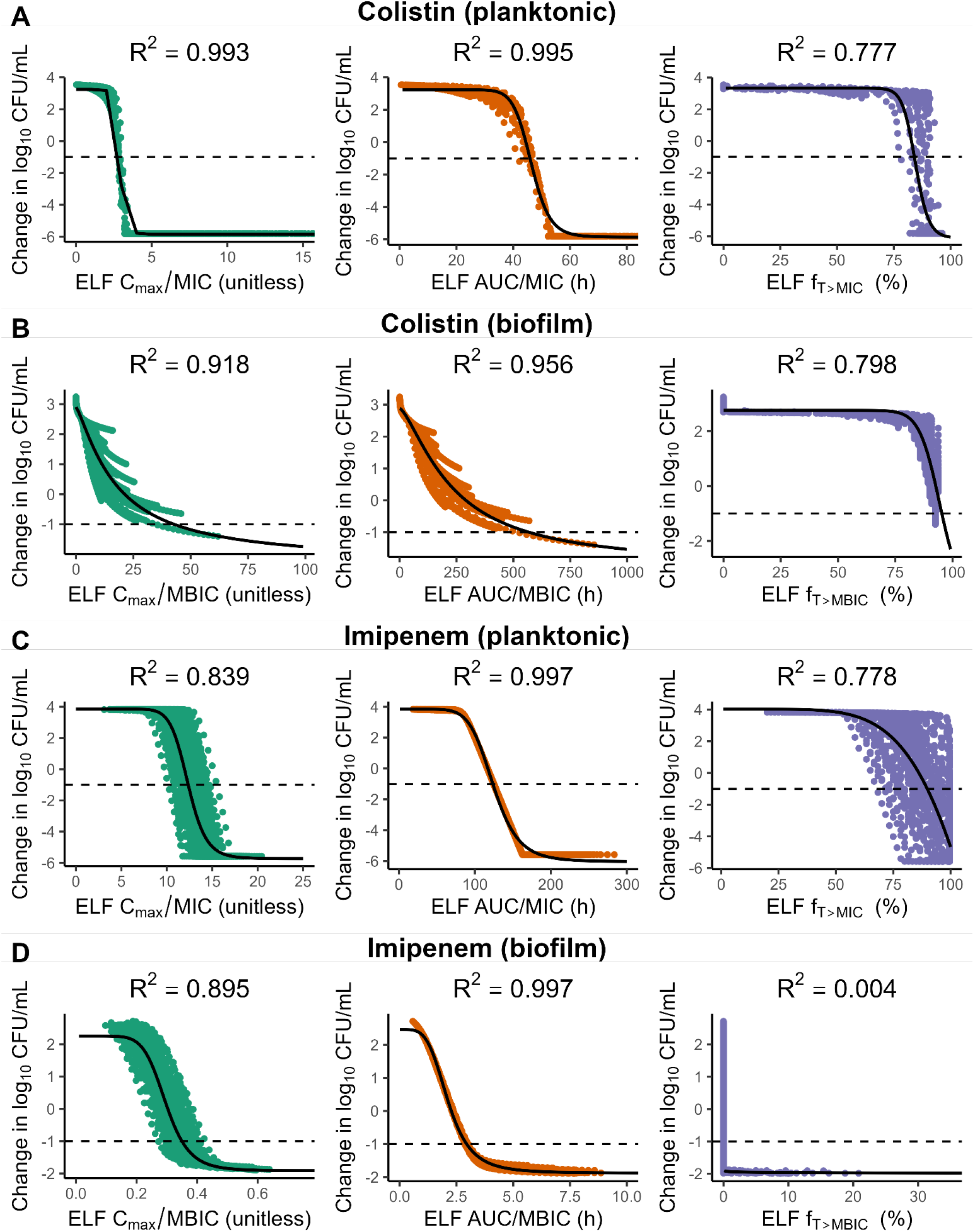
Pharmacokinetic-pharmacodynamic (PK-PD) target analysis for colistin and imipenem against planktonic and biofilm infections based on epithelial lining fluid (ELF) antibiotic concentrations. Dose fractionation studies were simulated and resulting ELF concentrations were regressed against the change in model predicted bacterial densities at 24 hours (points) compared to the baseline using a sigmoidal E_max_ model resulting in optimal model fits (solid lines).

In terms of PK-PD indices based on plasma drug concentration, AUC/MIC (R^2^=0.113) and AUC/MBIC (R^2^=0.171) were best correlated with the change of bacteria count for colistin, while AUC/MIC (R^2^=0.997) and AUC/MBIC (R^2^=0.997) were still most predictive of the effect for imipenem (**Figure S3**). A comparison between ELF-based (**Figure 6**) and plasma-based PK-PD indices (**Figure S3**) revealed an increased variation in responses to colistin and a consistent correlation between PK-PD indices and treatment effect for imipenem.

## DISCUSSION

We developed a pharmacodynamic model to characterize growth and kill dynamics from *in vitro* biofilm assays, focusing on the pathogen *P. aeruginosa* treated with colistin or imipenem, as a proof-of-concept. Prior knowledge of patient-specific antibiotic PK was integrated into the modeling framework and applied to investigate the effect of different dosing regimens, and the predictivity of different drug exposures (PK-PD indices) for bacterial response was assessed in clinical settings.

In this study, we derived models for different drugs, i.e., colistin and imipenem, as well as bacterial lifestyles, i.e., planktonic cells and biofilms. This model-based data-driven strategy enabled the evaluation of hypotheses with respect to differences in pharmacodynamic responses of planktonic versus biofilm cells. E_max_ models were identified for only two concentration-effect relationships while for the rest a simple linear model was identified, likely due to insufficient data availability, especially for high drug concentrations. The developed PD models facilitated the comparison of drug-, biological system- or experiment-specific parameters that ultimately characterize the observed response, providing insight into the underlying pharmacological basis of the antibiotic response. We identified distinct factors that contribute to increased resistance of biofilm cells compared to planktonic cells against both colistin and imipenem. First of all, biofilm cells exhibited slower growth rates (0.615 and 0.441 h^-1^) compared to planktonic cells (0.807 and 0.944 h^-1^), consistent with the literature [29, 30]. These reduced growth rates reflect biofilm cells’ adaptation to environments with limited nutrients and oxygen, which requires lower metabolic activity for survival. Secondly, a delay in antibiotic drug effect for biofilms, as represented by a transit model, was found for both colistin and imipenem against biofilm bacterial infections. This delay can be explained by the diffusion barrier presented by the extracellular matrix secreted by biofilm cells, the components of which could interact with antibiotics and slow their delivery. For example, the negatively charged polysaccharides could bind to positively charged colistin and impede penetration into biofilm [31]. Thirdly, compared to planktonic resistant species, a reduced susceptibility of biofilm resistant species was found (**Figure 3**). This discrepancy might be relevant to further physiological adaptations in biofilm cells compared to planktonic cells [32].

Coupling PD models with clinical population PK models enabled us to make translational predictions about the expected effects of bacterial growth/kill dynamics in patients. We predicted for colistin that dosing of 2 MIU per inhalation q 8 h shows a better biofilm-eradicating effect in lung infection patients compared to intravenous treatment of 2 MIU q 8 h and 9 MIU q 24 h. This result is in line with previous studies in COPD patients and ICU patients with pulmonary infections after lung transplantation [33, 34]. The regrowth of both planktonic and biofilm growing *P. aeruginosa* under exposure to colistin (planktonic: 1mg/L - 4 mg/L, biofilm: 1 mg/L - 64 mg/L) (**Figure 2A**) is probably due to the adaptive response that involved modifications of lipopolysaccharide (LPS) in the outer membrane, which prevents penetration of colistin [35–37]. Imipenem showed less predicted efficacy against planktonic and biofilm bacterial infections under standard clinical dosing, which might be partially because *P. aeruginosa* can readily develop adaptive responses (adaptive resistance) to imipenem and regrow under exposure to imipenem (**Figure 2B**) via upregulating production of alginate and AmpC β-lactamase [38, 39]. Although clinical implications of these model-based predictions should be treated with caution, they could provide additional insights into the differences observed in preclinical *in vivo* or *in vitro* studies, for instance when evaluating and comparing treatment regimens.

ELF concentration-based AUC/MIC and AUC/MBIC were identified as the PK-PD indices that could best predict the treatment outcome for both colistin and imipenem based on *in silico* dose fractionation simulations. A study using a murine infection model identified serum AUC/MIC and AUC/MBIC as the best PK-PD indices of colistin for planktonic and biofilm infections, yet for imipenem f_T>MIC_ and f_T>MBIC_ were found to best predict the efficacy for planktonic cell and biofilm infections [40]. Plasma samples are more readily measurable than lung ELF concentrations in patients, thus, we examined the feasibility of using plasma concentration for the derivation of PK-PD indices. Plasma concentration-based PK-PD indices of imipenem were found to be more predictive compared to those of colistin. For both colistin and imipenem, the same PK-PD indices (AUC/MIC and AUC/MBIC) were identified based on plasma concentration and ELF concentrations. Yet for colistin, indices (AUC/MIC and AUC/MBIC) identified with ELF concentrations (R^2^=0.995, 0.956) got less informative when replaced with plasma concentrations (R^2^=0.113, 0.171). This is because of colistin’s bi-directional transfer between plasma and ELF compartments in the population PK model [22], which leads to a non-linear relationship between plasma and ELF concentrations. The consistency for imipenem PKPD indices stems from its linear relationship between plasma and ELF concentration characterized in the population PK model [23]. Given that this speculation was based on simulations, further validation with real-world data is needed.

In the current study, pharmacodynamic models were developed based on bacteria data exposed to static concentrations of antibiotics measured up to 24 hours, which may not reflect the clinical reality for longer antibiotic treatment and time-varying antibiotic concentration. Conducting longer-term time-kill assays that mimic clinical treatments, such as using CDC biofilm reactors with humanized dynamic exposure, may help overcome the limitation [41, 42]. Indeed, several studies have employed CDC biofilm reactor model to investigate the efficacy of antibiotics against biofilms [30–32].

Our analysis demonstrates how a semi-mechanistic pharmacodynamic modeling approach can facilitate further pharmacological interpretation of *in vitro* biofilm infection models, through the separation of drug- and biological system (e.g. planktonic or biofilm)-specific parameters, in line with related publications describing similar semi-mechanistic mathematical models reported for *in vitro* flow cell models and an *in vivo* rat lung infection model [43–45]. We expect that semi-mechanistic modeling approaches are relevant to allow the simultaneous integration of multiple pertinent pharmacodynamic readouts, e.g. CFU counts, biofilm mass, and metabolic activity.

In conclusion, we report a proof-of-concept analysis of the utility of mathematical pharmacodynamic modeling of *in vitro* biofilm time-kill assays and its integration with clinical PK models to derive translational predictions about expected effects in patients.

## ACKNOWLEDGEMENTS

The research was financially supported by the China Scholarship Council (CSC).

## Notes

### Competing Interest Statement

The authors have declared no competing interest.

