## Supplementary materials for "Pharmacodynamic modeling of colistin and imipenem against *in vitro Pseudomonas aeruginosa* biofilms"

**Table S1**. Overview of experimental design information used for the *in vitro* experiments with planktonic cultures and the *in vitro* alginate bead biofilm experiments.

| **Antibiotic** | **Phenotype** | **Samples** | **Timepoints(h)** | **Concentrations (mg/L)** | **MIC (mg/L)** | **MBIC (mg/L)** |
| --- | --- | --- | --- | --- | --- | --- |
| **Colistin^#^** | Planktonic | 126 | 0,1,2,4,8,12,24 | 0,1,4,16,64,256 | 2 | - |
|  | Biofilm | 126 | 1,2,4,8,12,24 | 0,1,4,16,64,256 | - | 8 |
| **Imipenem** | Planktonic | 168 | 0,1,2,4,8,12,24 | 0,0.5,1,2,4,8,16,32 | 1 | - |
|  | Biofilm | 168 | 0,1,2,4,8,12,24 | 0,0.5,2,8,32,128,512,2048 | - | 32 |

^#^ The concentrations of colistin in time-kill studies were demonstrated in milligram per liter (mg/L). Colistimethate sodium (CMS), used as a prodrug in clinical settings, is commonly represented in million international units (MIU). Unit conversion:1 MIU CMS = 80 mg CMS.

Abbreviations: MIC, minimal inhibitory concentration; MBIC, minimal biofilm inhibitory concentration

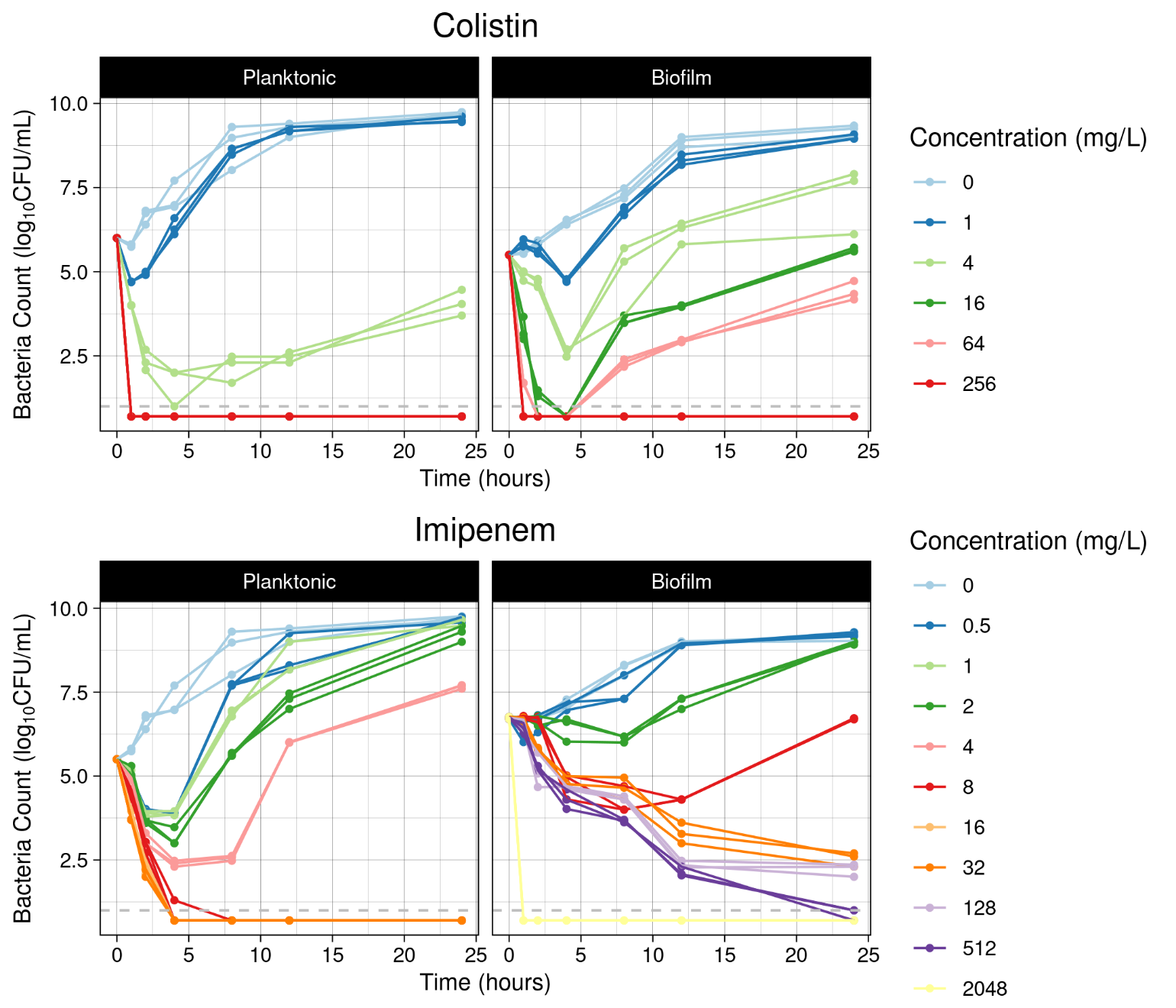

**Figure S1.** Raw observations for bacterial counts over time for colistin and imipenem in planktonic cultures and alginate-bead biofilm cultures. Observations below the quantification limit (gray dashed line) were displayed as half of the quantification limit for illustration.

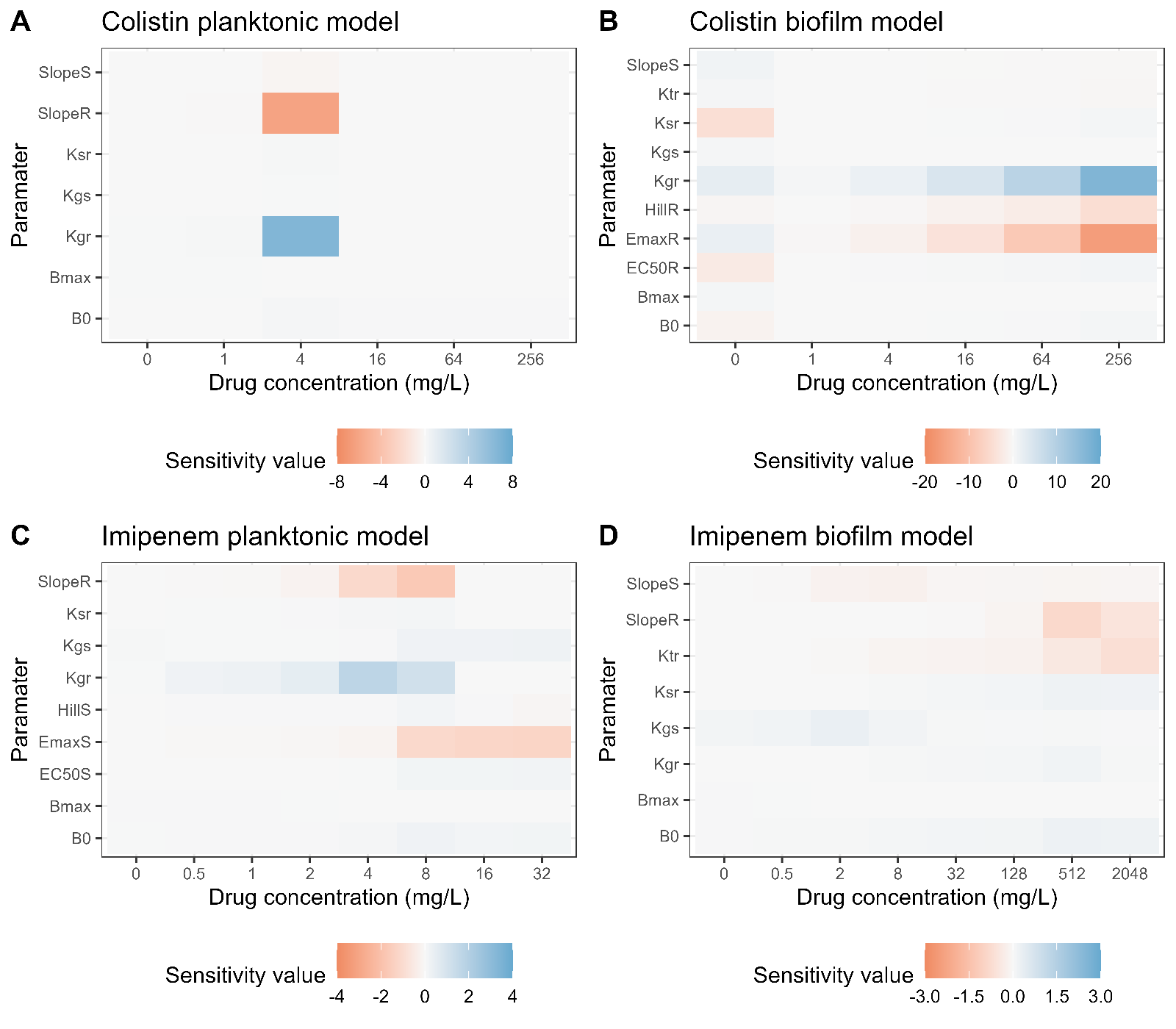

**Figure S2. Local parameter sensitivity analysis.** The colored tiles are the sensitivity value change of model output area under the bacterial growth curve (AUC_0-24h_) upon 10% increase in parameters for colistin planktonic model (A), colistin biofilm model (B), imipenem planktonic model(C) and imipenem biofilm model (D). The blue tiles represent a positive correlation between the parameter and model response, and the red tiles represent a negative correlation.

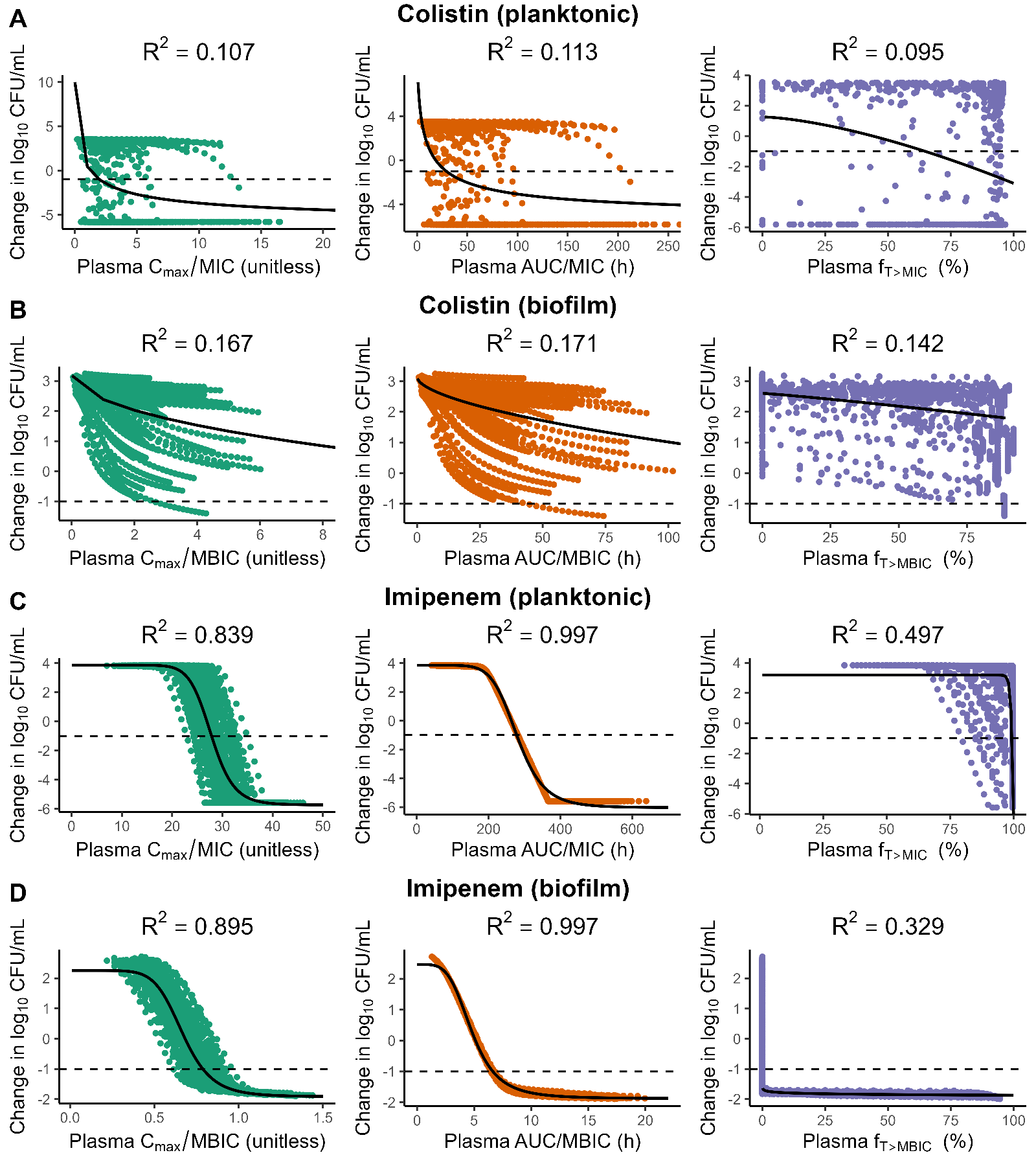

**Figure S3. Pharmacokinetic-pharmacodynamic (PK-PD) target analysis for colistin and imipenem against planktonic and biofilm infections based on plasma antibiotic concentrations.** Dose fractionation studies were simulated and resulting plasma concentrations were regressed against the change in model-predicted bacterial densities at 24 hours (points) compared to the baseline using a sigmoidal E_max_ model resulting in optimal model fits (solid lines).
